# Exo-enzymatic addition of diazirine-modified sialic acid to cell surfaces enables photocrosslinking of glycoproteins

**DOI:** 10.1101/2021.10.07.463072

**Authors:** Nageswari Yarravarapu, Rohit Sai Reddy Konada, Narek Darabedian, Nichole J. Pedowitz, Soumya N. Krishnamurthy, Matthew R. Pratt, Jennifer J. Kohler

**Affiliations:** Department of Biochemistry, UT Southwestern, Dallas, TX 75390 USA; Department of Chemistry, University of Southern California, Los Angeles, CA 90089 USA; Department of Biological Sciences, University of Southern California, Los Angeles, CA 90089 USA

## Abstract

Glycan binding often mediates extracellular macromolecular recognition events. Accurate characterization of these binding interactions can be difficult because of dissociation and scrambling that occur during purification and analysis steps. Use of photocrosslinking methods has been pursued to covalently capture glycan-dependent interactions *in situ* however use of metabolic glycan engineering methods to incorporate photocrosslinking sugar analogs is limited to certain cell types. Here we report an exo-enzymatic labeling method to add a diazirine-modified sialic acid (SiaDAz) to cell surface glycoconjugates. The method involves chemoenzymatic synthesis of diazirine-modified CMP-sialic acid (CMP-SiaDAz), followed by sialyltransferase-catalyzed addition of SiaDAz to desialylated cell surfaces. Cell surface SiaDAz-ylation is compatible with multiple cell types and is facilitated by endogenous extracellular sialyltransferase activity present in Daudi B cells. This method for extracellular addition of α2-6-linked SiaDAz enables UV-induced crosslinking of CD22, demonstrating the utility for covalent capture of glycan-mediated binding interactions.

## Introduction

A key challenge for glycoscience is the detection of glycan-mediated binding interactions within their native settings. Glycans mediate many physiologically critical recognition events; well-known examples include the recruitment of leukocytes to sites of inflammation through selectin-glycan interactions^1^ and attachment of influenza viral particles to mammalian cells through recognition of host glycans.^2^ Individual glycan-protein binding interactions are typically low-affinity, often exhibiting micromolar or even millimolar equilibrium dissociation constants. Notwithstanding these low affinities, glycan recognition events can be highly specific, often through multivalent presentation of glycans and/or their ligands on cell surfaces.^3-4^ Typical biochemical methods for interaction discovery break apart cells, resulting in disruption of and scrambling of glycan-protein interactions. To solve this challenge, we and others have used metabolic glycan engineering approaches to introduce photoactivatable crosslinking groups (photocrosslinkers) onto cellular glycans.^5-10^ These strategies involve preparing monosaccharide analogs functionalized with photocrosslinkers. Such photocrosslinking sugars can be metabolized by cells and incorporated into glycoconjugates in place of normal monosaccharides. Subsequent activation of the photocrosslinker by application of UV irradiation results in formation of a covalent crosslink between the glycoconjugate and nearby molecules. Once this crosslink is formed, then cells can be lysed and the covalent complexes analyzed by common methods such as immunoblot and mass spectrometry-based proteomics.

Sialic acids are a class of nonulosonic acids found at the non-reducing termini of many cell surface glycans. By virtue of their position at the termini of glycoconjugates, unusual 9-carbon structure, and negative charge, sialic acids mediate many extracellular recognition events.^11^ N-acetylneuraminic acid (Neu5Ac) is the most common sialic acid in human cells and is biosynthesized from N-acetylmannosamine (ManNAc). To enable introduction of a photocrosslinker onto sialic acid residues, we previously reported a cell-permeable analog of ManNAc in which the diazirine photocrosslinker is appended to the N-acyl sidechain and the polar hydroxyl groups are masked with biolabile acetyl groups (Ac_4_ManNDAz).^7, 12^ We observed metabolism of Ac_4_ManNDAz to its sialic acid counterpart (SiaDAz), which can be incorporated into both glycolipids and glycoproteins in place of naturally occurring sialic acid.^13-16^ The diazirine photocrosslinker is activated by UV irradiation, resulting in covalent crosslinking of glycan-mediated interactions. These include the sialic acid-dependent dimerization of the sialic acid-binding immunoglobulin-type lectin (Siglec) CD22,^7, 12-13, 17^ recognition of the sialylated glycolipid (ganglioside) GM1 by cholera toxin,^14, 18^ and interactions between a glycan-recognizing antibody and cell surface glycoproteins.^19^ While SiaDAz is a potentially powerful tool, we have encountered limitations in its use. Most notably, cell types vary in their ability to metabolize Ac_4_ManNDAz to cell surface SiaDAz. For example, in Jurkat cells, Ac_4_ManNDAz is metabolized efficiently resulting in 65 % or more of the cell surface sialic acid being diazirine-modified.^13, 16^ However, when K562, HeLa, or PC-3 cells were cultured with Ac_4_ManNDAz, no cell surface SiaDAz was detected.^16^ Use of alternative precursors did not solve cell type-specific metabolic limitations.^16^ A second limitation in the metabolic engineering approach lies in the lack of specificity of incorporation. SiaDAz is added by both α2-6- and α2-3-sialyltransferases without targeting to specific glycoconjugates of interest.^13^

To overcome limitations imposed by the metabolic incorporation strategy, we decided to use recombinant enzymes to add SiaDAz to cell surface glycoconjugates. Our approach is inspired by prior studies that used neuraminidases (sialidases) and sialyltransferases to remodel the cell surface sialome.^20^ Most directly relevant to our goal, Boons, Steet, and co-workers employed a selective exo-enzymatic labeling (SEEL) strategy to add azide-modified or biotinylated sialic acid to cell surface glycoconjugates.^21-23^ These authors prepared modified CMP-sialic acid analogs chemoenzymatically, then used mammalian sialyltransferases ST6GAL1 and ST3GAL1 to transfer the sialic acid analogs to cell surface glycoproteins

Here we report the preparative-scale chemoenzymatic synthesis of diazirine-modified CMP-sialic acid (CMP-SiaDAz). We show that *Photobacterium damsela* α2-6-sialyltransferase can transfer SiaDAz from CMP-SiaDAz to the surface of cells that have been desialylated by *Arthrobacter ureafaciens* neuraminidase. Cell surface SiaDAz-ylation can be performed in multiple cell types, including three cell lines in which metabolic incorporation of SiaDAz was unsuccessful. Notably, inclusion of *P. damsela* α2-6-sialyltransferase is not required for addition of α2-6-linked SiaDAz to Daudi B cells, suggesting that these cells harbor an endogenous extracellular α2-6-sialyltransferase activity. Finally, we report that extracellular addition of α2-6-linked SiaDAz enables UV-induced crosslinking of the CD22 Siglec, demonstrating the utility of method.

## Results and Discussion

To support photocrosslinking applications, our goal was to replace normal cell surface sialic acids, such as Neu5Ac, with SiaDAz, a photocrosslinking sialic acid analog (**Fig. 1a**). To accomplish this, we envisioned removing Neu5Ac a linkage-agnostic manner using a non-selective neuraminidase. In a subsequent step, SiaDAz would be added exclusively in the α2-6 linkage using a sialyltransferase. The strategy requires purified neuraminidase and sialyltransferase, both of which are commercially available, as well as the unnatural nucleotide-sugar CMP-SiaDAz. We prepared CMP-SiaDAz through a chemoenzymatic approach, similar to methods reported for other CMP-sialic acids (**Fig. 1b**).^24-27^ Diazirine-modified ManNAc (ManNDAz) was prepared by chemical synthesis, as described previously.^7, 16^ ManNAc, pyruvate, and CTP were incubated together with microbial N-acetylneuraminic acid aldolase and *Neisseria meningitidis* CMP-sialic acid synthetase (NmCSS) in the presence of Mg^2+^. Alkaline phosphatase was included promote product formation. Consistent with our prior results,^28^ both the aldolase and NmCSS accepted substrates with the diazirine modification. CMP-SiaDAz produced in this reaction was purified by Bio-Gel P-2 column chromatography. The identity of CMP-SiaDAz was established by ^1^H, ^13^C, and ^31^P NMR (**Supplemental Figures 1-3**) and high-resolution mass spectrometry (HRMS); purity was evaluated by HPLC (**Supplemental Figure 4**).

**Figure 1.**
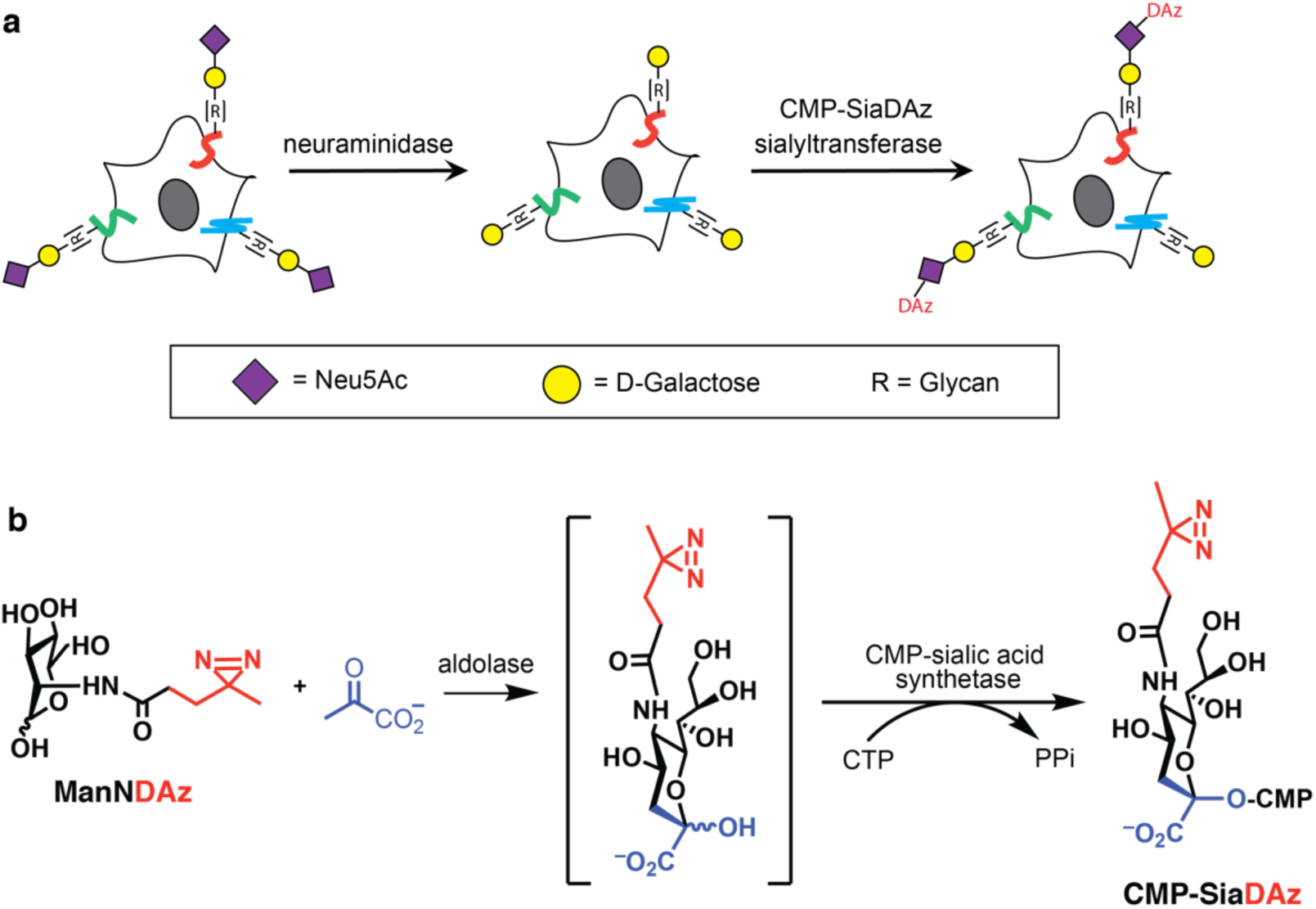
Exo-enzymatic labeling strategy to add photocrosslinking sialic acid (SiaDAz) to cell surface glycoconjugates. (**a**) Existing cell surface sialic acids are removed by a neuraminidase, followed by sialyltransferase-catalyzed addition of α2-6-linked SiaDAz. (**b**) One-pot chemoenzymatic synthesis of CMP-SiaDAz for use in exo-enzymatic labeling.

We identified conditions to enzymatically add SiaDAz to cell surface glycoconjugates. We began with Jurkat cells, a cell line where we have previously introduced SiaDAz through metabolic glycan engineering.^13, 16, 29^ Jurkat cell surfaces were treated with *A. ureafaciens* neuraminidase, then cell surface sialic acid was detected by flow cytometry using *Sambucus nigra* agglutinin (SNA; α2-6 sialic acids) or *Maackia amurensis* lectin II (MAL II; α2-3 sialic acids). *A. ureafaciens* neuraminidase efficiently removed both α2-6- and α2-3-linked sialic acids (**Fig. 2a** and **Supplemental Figure 5a**) and subsequent treatment with *P. damsela* α2-6-sialyltransferase and CMP-Neu5Ac restored cell surface α2-6 sialylation to a level similar to that observed in untreated cells (**Fig. 2a and b**). Similarly, when neuraminidase-treated cells were incubated with *P. damsela* α2-6-sialyltransferase and CMP-SiaDAz, increased SNA binding was observed, consistent with the addition of SiaDAz in an α2-6 linkage.

**Figure 2.**
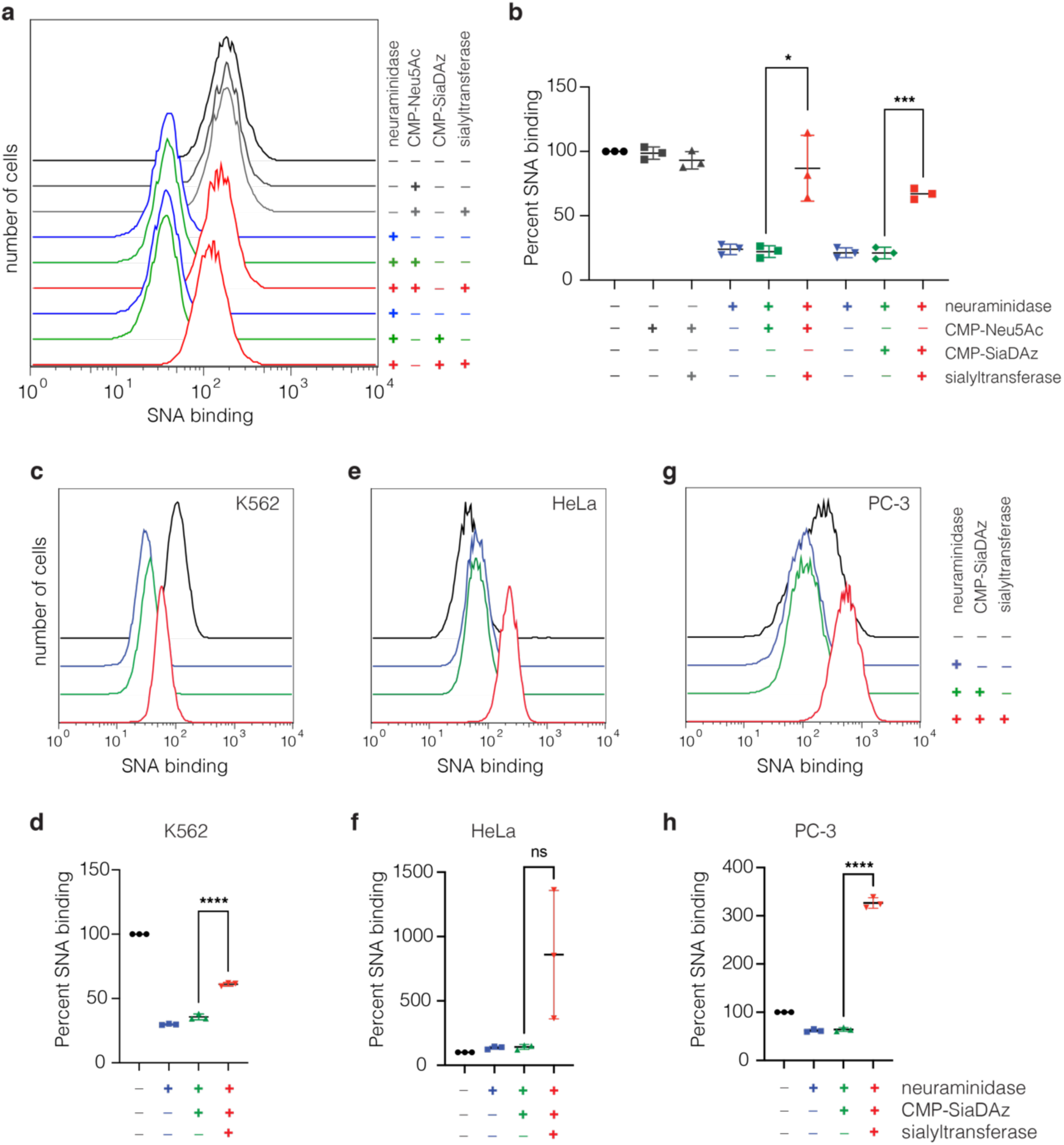
Enzymatic addition of Neu5Ac or SiaDAz to cell surface glycoconjugates. (**a**) Jurkat cells were treated with or without *Arthrobacter ureafaciens* neuraminidase, followed by treatment with *Photobacterium damsela* α2-6-sialyltransferase in the presence of CMP-Neu5Ac, CMP-SiaDAz, or no nucleotide sugar. Cell surface α2−6-linked sialic acids were detected by flow cytometry using *Sambucus nigra* agglutinin (SNA). (**b**) Quantification of flow cytometry analyses performed as in panel **a**. Mean fluorescence intensity (MFI) of each sample was calculated and normalized to that measured for cells not treated with neuraminidase or sialyltransferase. Error bars represent the standard deviation of two biological replicates. (**c, e, g**) K562, Hela, and PC-3 cells were treated as in panel a, and cell surface α2−6-linked sialic acids were detected by flow cytometry using SNA. (**d, f, h**) Quantification of flow cytometry analyses performed as in panel c. Mean fluorescence intensity (MFI) of each sample was calculated and normalized to that measured for cells not treated with neuraminidase or sialyltransferase. Error bars represent the standard deviation of three biological replicates. Statistical significance was assessed by unpaired t-test with * p < 0.05; *** p < 0.001; **** p < 0.0001.

Next, we evaluated whether exo-enzymatic addition of SiaDAz could be performed in additional cell types. We selected three cell lines (K562, HeLa, and PC-3) in which we had previously been unable to observe cell surface SiaDAz after a metabolic labeling protocol in which the cells were cultured with the Ac_4_ManNDAz precursor.^16^ In K562 cells, *A. ureafaciens* neuraminidase efficiently removed endogenous α2-6 sialic acids, and treatment with *P. damsela* α2-6-sialyltransferase and CMP-SiaDAz restored SNA binding although not to the level observed in untreated cells (**Fig. 2c and d, Supplemental Figure 5b**). In contrast, *A. ureafaciens* neuraminidase treatment had no effect on SNA binding to HeLa cells, but treatment with *P. damsela* α2-6-sialyltransferase and CMP-SiaDAz resulted in variably increased SNA binding (**Fig. 2e and f, Supplemental Figure 5c**). Finally, for PC-3 cells, neuraminidase treatment removed endogenous α2-6 sialic acids, but the amount of SiaDAz added by *P. damsela* α2-6-sialyltransferase greatly exceeded the initial level of α2-6 sialylation (**Fig. 2g and h, Supplemental Figure 5d**). We speculate that the inability of *A. ureafaciens* neuraminidase treatment to affect SNA binding to HeLa cells may reflect a low level of endogenous α2-6-sialylation in these cells;^30^ similarly, PC-3 cells may display terminal Gal residues that are normally unsialylated but are substrates for SiaDAz-ylation. Taken together, these results demonstrate that exo-enzymatic addition of SiaDAz can be performed in multiple cell types although the SiaDAz added by *P. damsela* α2-6-sialyltransferase may not faithfully reproduce the normal sialylation pattern of the cells.

We also used exo-enzymatic labeling to add α2-6-linked SiaDAz to the surface of Daudi B cells. Similar to K562 cells, we found that neuraminidase treatment effectively reduced cell surface α2-6 sialylation and subsequent treatment with *P. damsela* α2-6-sialyltransferase and CMP-SiaDAz led to increased SNA binding consistent with the addition of α2-6-linked SiaDAz (**Fig. 3a and b**). However, we were surprised to observe increased SNA binding when neuraminidase-treated cells were incubated with CMP-SiaDAz in the absence of exogenously-added sialyltransferase. This result suggests that Daudi cells endogenously produce extracellular α2-6-linked sialyltransferase activity, as has been reported previously for human and mouse B cells.^31-32^

**Figure 3.**
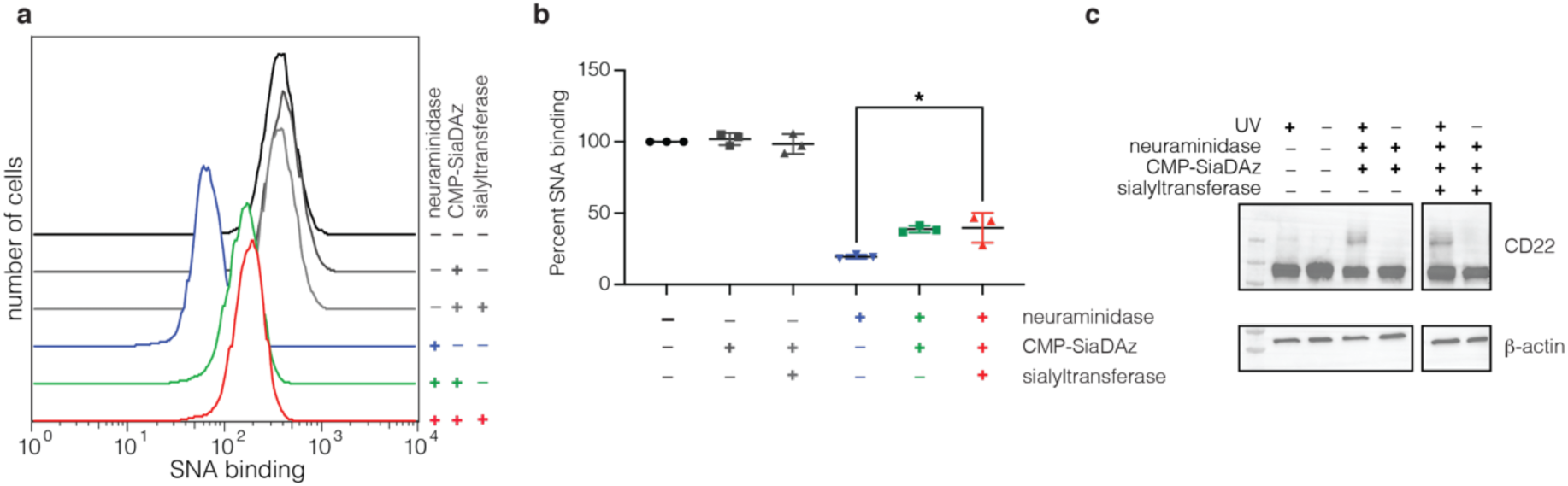
Exo-enzymatically added SiaDAz mediates photocrosslinking of CD22 oligomers in Daudi cells. (**a**) Daudi cells were treated with or without *A. ureafaciens* neuraminidase, followed by treatment with *P. damsela* α2-6-sialyltransferase in the presence of CMP-SiaDAz or no nucleotide sugar. Cell surface α2−6-linked sialic acids were detected by flow cytometry using SNA. (**b**) Quantification of analyses performed as in panel **a**. Mean fluorescence intensity (MFI) of each sample was calculated and normalized to that measured for cells not treated with neuraminidase or sialyltransferase. Error bars represent the standard deviation of three biological replicates. (**c**) Daudi cells were treated with *A. ureafaciens* followed by exo-enzymatic labeling with *P. damsela* α2-6-sialyltransferase in the presence or absence of CMP-SiaDAz. After labeling, cells were subjected to UV irradiation, followed by lysis. Cell contents were analyzed by immunoblot, using an anti-CD22 antibody. Data presented are representative of three trials. Statistical significance was assessed by unpaired t-test with * p < 0.05.

Finally, we tested whether exo-enzymatically added SiaDAz could be used to crosslink sialylated proteins to their binding partners. As a test case, we selected the sialic acid-dependent oligomerization of the Siglec CD22. CD22 contains an extracellular V-set domain responsible for recognition of sialosides and is itself modified with N-linked glycans. Covalent capture of CD22 oligomers has previously been achieved using metabolically incorporated photocrosslinking sialic acid analogs.^5, 7^ Daudi cells were treated with neuraminidase followed by exo-enzymatic addition of SiaDAz. SiaDAz-ylated cells were subjected to 365 nm irradiation for 10 min, then immediately collected and lysed. Cell lysates were analyzed by immunoblot using an anti-CD22 antibody. In lysates from untreated cells, CD22 exhibited an apparent molecular weight of ∼130 kDa. In lysates from cells in which crosslinking was performed, a second band was detected with apparent molecular weight consistent with formation of a CD22 dimer (**Fig. 3c**). Appearance of this crosslinked band was dependent on inclusion of CMP-SiaDAz but not *P. damsela* α2-6-sialyltransferase, suggested that the endogenous sialyltransferase activity of Daudi cells was sufficient to support the addition of SiaDAz for crosslinking. The small amount of background crosslinking detected in cells that were UV irradiated but not SiaDAz-ylated may be due to UV-induced formation of free radicals.^33^

In sum, the results presented here expand the scope of photocrosslinking sugar technology as well as the information that can be obtained using it. Specifically, using the exo-enzymatic labeling method, SiaDAz can be added to cell surface glycoconjugates from any cell line. Also, since exo-enzymatic SiaDAz addition is linkage-specific, the method also provides information about the sialic acid linkage needed to mediate a specific binding interaction. Use of a bacterial sialyltransferase makes the method accessible, since these enzymes are simple to express and purify and are often commercially available. However, one limitation is that bacterial sialyltransferases may exhibit different substrate specificities than their mammalian counterparts. In the case of Daudi cells, we were able to take advantage of the cell’s endogenous extracellular α2-6-sialyltransferase activity, likely due to ST6GAL1. Therefore, in this case, the α2-6-SiaDAz-ylation pattern would be expected faithfully mimic to the normal α2-6-sialylation of these cells. Notably, we observed SiaDAz-mediated CD22 crosslinking in Daudi cells, indicating the normal cell surface interactions can occur after the exo-enzymatic treatment. Potential uses of this method include characterization of interactions occurring among cell surface glycoproteins, as well as identification of the cell surface receptors for bacterial and viral lectins.

## Acknowledgment

The authors thank Dr. Vaishnavi Nair for experimental assistance. Funding was provided by the Welch Foundation (I-1686) and the NIH (R01GM125939).

## METHODS

### Materials

N-Acetylneuraminic acid aldolase (cat # NAL-301) was purchased from Toyobo. CMP-sialic acid synthetase from *Neisseria meningitidis* group B (NmCSS, cat # C1999), neuraminidase (sialidase) from *Arthrobacter ureafaciens* (cat # 10269611001) and α2-6-sialyltransferase from *Photobacterium damsela* (Pd2-6ST, cat # S2076) were purchased from Sigma. CMP-Neu5Ac (cat # 233264) was purchased from Millipore. Biotinylated *Sambucus nigra* agglutinin (SNA, cat # B-1305) was purchased from Vector Labs. Alkaline phosphatase (cat # M0290S) was purchased from NEB. DTAF-streptavidin (cat # 016-010-084) was purchased from Jackson Immunoresearch. Anti-rabbit CD22 monoclonal antibody (cat # EP498Y) and anti-actin antibody (cat # Ab8227) were purchased from Abcam. Anti-rabbit antibody conjugated to HRP (cat # 65-6120) was purchased from Invitrogen. SuperSignal™ West Pico PLUS Chemiluminescent Substrate (cat # 34580) was purchased from Thermo Scientific.

### Chemoenzymatic synthesis of CMP-SiaDAz

Synthesis of ManNDAz has been reported previously.^7, 16^ CMP-SiaDAz was synthesized according to previously published method (PMID: 15556760). Pilot, small-scale (50 μl reaction mixture volume) synthesis of CMP-SiaDAz was performed in 100 mM Tris–HCl pH 8.8 containing 20 mM MgCl_2_, 10 mM ManNDAz, 50 mM sodium pyruvate, and 10 mM CTP. To the reaction mixture, 10 mU N-acetylneuraminic acid aldolase and 10 mU CMP–sialic acid synthetase were added. The reaction mixture was incubated for 6 h at 37 °C then quenched by addition of equal volume of MeOH. Precipitate was removed by centrifugation and then filtration (0.2 μm syringe filter). Aliquots of 10 μL were injected for HPLC analysis. Preparative synthesis of CMP–SiaDAz (25 mL reaction mixture volume) was performed in 100 mM Tris–HCl buffer pH 8.8 containing 50 mg ManNDAz, 20 mM MgCl_2_, 5 equivalents sodium pyruvate, 1 equivalent CTP, 6 U N-acetylneuraminic acid aldolase, and 6 U CMP–sialic acid synthetase. The reaction mixture was incubated at 37 °C overnight in an incubator with shaking (140 rpm). The reaction was stopped by addition of equal volume of MeOH. Precipitate was removed by centrifugation. The supernatant was dried by rotary evaporation and resuspended in 2.5 mL of water. For purification, 0.5ml of concentrated reaction mixture was loaded on BioGel P-2 gel filtration column (1.0 × 100 cm) and 0.5 mL fractions were collected using water as elution buffer. Characterization data are presented in the **Supplementary Figures 1-4**.

### Cell culture

All cell lines were cultured at 37 °C, 5 % CO2 in a water saturated environment. K562 and Jurkat cells were obtained from ATCC. Daudi cells were obtained from Ellen Vitetta (UT Southwestern Medical Center). K562, Jurkat and Daudi cells were cultured in RPMI 1640 medium containing 2 mM glutamine, 10 % FBS, and 1 % Pen/Strep. HeLa cells were obtained from the ATCC and were cultured in DMEM containing 4.5 g/L D-glucose, 110 mg/L sodium pyruvate, 4 mM L-glutamine, 10 % fetal bovine serum (FBS), and 1 % Pen/Strep. PC-3 cells were obtained from Jer-Tsong Hsieh (UT Southwestern) and cultured in DMEM containing 4.5 g/L D-glucose, 110 mg/L sodium pyruvate, 4 mM L-glutamine, 10 % FBS, and 1 % Pen/Strep.

### Exo-enzymatic labeling

For enzymatic labeling, 3 × 10^6^ cells were used for each sample. Cells were washed three times with 1 mL Dulbecco’s phosphate buffered saline (DPBS) and incubated in 300 μL of serum-free culture medium containing BSA (13.3 μg/mL) and neuraminidase (50 mM) for 2 h at 37 °C. Cells were washed three times with 1 mL DPBS, then incubated for 30 min at 37 °C in 300 μL of serum-free culture medium containing BSA, 1 mM CMP-SiaDAz, 0.35U Pd2-6ST, and 2 μL alkaline phosphatase (10U/μL). Cells were washed three times with 1 mL DPBS before flow cytometry analysis or crosslinking experiments.

### Flow cytometry analysis

For detection of α2-6-linked sialic acids and α2-3-linked sialic acids, 4 × 10^6^ cells were incubated for 30 min on ice with 100 μL of 2.67 μg/mL SNA-biotin or 10 μg/mL MAL II-biotin, respectively. Cells were washed twice in 200 μL ice-cold phosphate-buffered saline (PBS) containing 0.1 % (w/v) BSA. Then cells were incubated with 100 μL of 7.7 μg/mL DTAF-streptavidin for 30 min on ice in the dark. Cells were washed three times in 200 μL PBS containing 0.1 % (w/v) BSA, then resuspended in 500 μL PBS containing 0.1 % (w/v) BSA and 1.25 μg/mL propidium iodide. Samples were analyzed on a FACS Calibur on the FL1 and FL3 channels. Cells were gated based on forward scatter versus side scatter, and cells with high PI staining were excluded. For each sample, 10,000 live cell events were recorded.

### Crosslinking

After exo-enzymatic labeling, Daudi cells were resuspended in DPBS, transferred to 6-well plate placed on ice, then irradiated with UV light (365 nm, UVP, XX-20BLB lamp) for 10 min. Control cells were treated in the same way but not irradiated. Cells were lysed by resuspending the cell pellet in 150 μl of lysis buffer (300 mM NaCl, 50 mM Tris-Cl pH 8, 1% (vol/vol) Triton X-100, 1 mM EDTA pH 8, SIGMAFAST™ protease inhibitor cocktail cat # S8830) and incubating on ice for 1 h. Cell lysates were centrifuged at 14,000*g* for 4 min at room temperature. The supernatant was transferred to a clean microcentrifuge tube. For immunoblot analysis, samples were loaded with Laemmli sample buffer containing 50 mM DTT and proteins were resolved on 7.5% SDS-PAGE then transferred to PVDF membranes. Membranes were blocked with 5 % (w/v) non-fat milk for 1 h, then incubated with 1:2000 dilution of anti-rabbit CD22 monoclonal antibody (Abcam: EP498Y) in 25 mM Tris buffered saline containing 0.1% Tween (TBST, pH 7.4) containing 5 % (w/v) non-fat milk overnight at 4 °C. Membranes were washed three times in TBST and further incubated with 1:2000 Goat anti-Rabbit IgG-HRP conjugated antibody for 1 h at room temperature. Washing step was repeated with TBST and the membranes were developed using SuperSignal™ West Pico PLUS Chemiluminescent Substrate, imaged on a ChemiDocTM MP Imaging System (Bio-Rad). For actin loading control, membranes were blocked in 5 % (w/v) NFM PBST for 1 hr at room temperature. Membranes were then incubated with an anti-actin antibody (1:2000, Abcam cat # Ab8227) for overnight at 4°C in PBST containing 5 % (w/v) NFM. Membranes were washed and then incubated with a goat anti-rabbit antibody conjugated to HRP (1:2000, Invitrogen cat # 65-6120) in PBST containing 5 % (w/v) non-fat milk. After washing, membranes were developed using SuperSignal™ West Pico PLUS Chemiluminescent Substrate, imaged on a ChemiDocTM MP Imaging System (Bio-Rad).

## References

1. Lowe, J. B., Glycan-dependent leukocyte adhesion and recruitment in inflammation. Curr. Opin. Cell Biol. 2003, 15 (5), 531–8.

2. Thompson, A. J.; de Vries, R. P.; Paulson, J. C., Virus recognition of glycan receptors. Curr Opin Virol 2019, 34, 117–129.

3. Collins, B. E.; Paulson, J. C., Cell surface biology mediated by low affinity multivalent protein-glycan interactions. Curr Opin Chem Biol 2004, 8 (6), 617–25.

4. Cairo, C. W.; Gestwicki, J. E.; Kanai, M.; Kiessling, L. L., Control of multivalent interactions by binding epitope density. J Am Chem Soc 2002, 124 (8), 1615–9.

5. Han, S.; Collins, B. E.; Bengtson, P.; Paulson, J. C., Homomultimeric complexes of CD22 in B cells revealed by protein-glycan cross-linking. Nat Chem Biol 2005, 1 (2), 93–7.

6. Luchansky, S. J.; Goon, S.; Bertozzi, C. R., Expanding the diversity of unnatural cell-surface sialic acids. Chembiochem 2004, 5 (3), 371–4.

7. Tanaka, Y.; Kohler, J. J., Photoactivatable crosslinking sugars for capturing glycoprotein interactions. J Am Chem Soc 2008, 130 (11), 3278–9.

8. Yu, S. H.; Boyce, M.; Wands, A. M.; Bond, M. R.; Bertozzi, C. R.; Kohler, J. J., Metabolic labeling enables selective photocrosslinking of O-GlcNAc-modified proteins to their binding partners. Proc Natl Acad Sci U S A 2012, 109 (13), 4834–9.

9. Feng, L.; Hong, S.; Rong, J.; You, Q.; Dai, P.; Huang, R.; Tan, Y.; Hong, W.; Xie, C.; Zhao, J.; Chen, X., Bifunctional unnatural sialic acids for dual metabolic labeling of cell-surface sialylated glycans. J Am Chem Soc 2013, 135 (25), 9244–7.

10. Wu, H.; Shajahan, A.; Yang, J. Y.; Capota, E.; Wands, A. M.; Arthur, C. M.; Stowell, S. R.; Moremen, K. W.; Azadi, P.; Kohler, J. J., A photo-cross-linking GlcNAc analog enables covalent capture of N-linked glycoprotein-binding partners on the cell surface. Cell Chem Biol 2021.

11. Chen, X.; Varki, A., Advances in the biology and chemistry of sialic acids. Acs Chem Biol 2010, 5 (2), 163–76.

12. Bond, M. R.; Zhang, H.; Vu, P. D.; Kohler, J. J., Photocrosslinking of glycoconjugates using metabolically incorporated diazirine-containing sugars. Nat Protoc 2009, 4 (7), 1044–63.

13. Bond, M. R.; Zhang, H.; Kim, J.; Yu, S. H.; Yang, F.; Patrie, S. M.; Kohler, J. J., Metabolism of diazirine-modified N-acetylmannosamine analogues to photo-cross-linking sialosides. Bioconjug Chem 2011, 22 (9), 1811–23.

14. Bond, M. R.; Whitman, C. M.; Kohler, J. J., Metabolically incorporated photocrosslinking sialic acid covalently captures a ganglioside-protein complex. Mol Biosyst 2010, 6 (10), 1796–9.

15. Whitman, C. M.; Yang, F.; Kohler, J. J., Modified GM3 gangliosides produced by metabolic oligosaccharide engineering. Bioorg Med Chem Lett 2011, 21 (17), 5006–10.

16. Pham, N. D.; Fermaintt, C. S.; Rodriguez, A. C.; McCombs, J. E.; Nischan, N.; Kohler, J. J., Cellular metabolism of unnatural sialic acid precursors. Glycoconj J 2015, 32 (7), 515–29.

17. Tanaka, Y.; Bond, M. R.; Kohler, J. J., Photocrosslinkers illuminate interactions in living cells. Mol Biosyst 2008, 4 (6), 473–80.

18. Wands, A. M.; Fujita, A.; McCombs, J. E.; Cervin, J.; Dedic, B.; Rodriguez, A. C.; Nischan, N.; Bond, M. R.; Mettlen, M.; Trudgian, D. C.; Lemoff, A.; Quiding-Jarbrink, M.; Gustavsson, B.; Steentoft, C.; Clausen, H.; Mirzaei, H.; Teneberg, S.; Yrlid, U.; Kohler, J. J., Fucosylation and protein glycosylation create functional receptors for cholera toxin. Elife 2015, 4, e09545.

19. McCombs, J. E.; Zou, C.; Parker, R. B.; Cairo, C. W.; Kohler, J. J., Enhanced Cross-Linking of Diazirine-Modified Sialylated Glycoproteins Enabled through Profiling of Sialidase Specificities. Acs Chem Biol 2016, 11 (1), 185–92.

20. Paulson, J. C.; Sadler, J. E.; Hill, R. L., Restoration of specific myxovirus receptors to asialoerythrocytes by incorporation of sialic acid with pure sialyltransferases. J Biol Chem 1979, 254 (6), 2120–4.

21. Mbua, N. E.; Li, X.; Flanagan-Steet, H. R.; Meng, L.; Aoki, K.; Moremen, K. W.; Wolfert, M. A.; Steet, R.; Boons, G. J., Selective exo-enzymatic labeling of N-glycans on the surface of living cells by recombinant ST6Gal I. Angew Chem Int Ed Engl 2013, 52 (49), 13012–5.

22. Sun, T.; Yu, S. H.; Zhao, P.; Meng, L.; Moremen, K. W.; Wells, L.; Steet, R.; Boons, G. J., One-Step Selective Exoenzymatic Labeling (SEEL) Strategy for the Biotinylation and Identification of Glycoproteins of Living Cells. J Am Chem Soc 2016, 138 (36), 11575–11582.

23. Yu, S. H.; Zhao, P.; Sun, T.; Gao, Z.; Moremen, K. W.; Boons, G. J.; Wells, L.; Steet, R., Selective Exo-Enzymatic Labeling Detects Increased Cell Surface Sialoglycoprotein Expression upon Megakaryocytic Differentiation. J Biol Chem 2016, 291 (8), 3982–9.

24. Blixt, O.; Paulson, J. C., Biocatalytic preparation of N-glycolylneuraminic acid, deaminoneuraminic acid (KDN) and 9-azido-9-deoxysialic acid oligosaccharides. Adv Synth Catal 2003, 345 (6-7), 687–690.

25. Yu, H.; Yu, H.; Karpel, R.; Chen, X., Chemoenzymatic synthesis of CMP-sialic acid derivatives by a one-pot two-enzyme system: comparison of substrate flexibility of three microbial CMP-sialic acid synthetases. Bioorgan Med Chem 2004, 12 (24), 6427–6435.

26. Yu, H.; Chokhawala, H. A.; Huang, S. S.; Chen, X., One-pot three-enzyme chemoenzymatic approach to the synthesis of sialosides containing natural and non-natural functionalities. Nature Protocols 2006, 1 (5), 2485–2492.

27. Gilormini, P. A.; Lion, C.; Noel, M.; Krzewinski-Recchi, M. A.; Harduin-Lepers, A.; Guerardel, Y.; Biot, C., Improved workflow for the efficient preparation of ready to use CMP-activated sialic acids. Glycobiology 2016, 26 (11), 1151–1156.

28. McCombs, J. E.; Zou, C. X.; Parker, R. B.; Cairo, C. W.; Kohler, J. J., Enhanced Cross-Linking of Diazirine-Modified Sialylated Glycoproteins Enabled through Profiling of Sialidase Specificities. Acs Chem Biol 2016, 11 (1), 185–192.

29. Bond, M. R.; Whitman, C. M.; Kohler, J. J., Metabolically incorporated photocrosslinking sialic acid covalently captures a ganglioside-protein complex. Molecular Biosystems 2010, 6 (10), 1796–1799.

30. Stamenkovic, I.; Sgroi, D.; Aruffo, A.; Sy, M. S.; Anderson, T., The B lymphocyte adhesion molecule CD22 interacts with leukocyte common antigen CD45RO on T cells and alpha 2-6 sialyltransferase, CD75, on B cells. Cell 1991, 66 (6), 1133–44.

31. Irons, E. E.; Lee-Sundlov, M. M.; Zhu, Y.; Neelamegham, S.; Hoffmeister, K. M.; Lau, J. T., B cells suppress medullary granulopoiesis by an extracellular glycosylation-dependent mechanism. Elife 2019, 8.

32. Irons, E. E.; Punch, P. R.; Lau, J. T. Y., Blood-Borne ST6GAL1 Regulates Immunoglobulin Production in B Cells. Front Immunol 2020, 11, 617.

33. Russ, M.; Lou, D.; Kohler, H., Photo-activated affinity-site cross-linking of antibodies using tryptophan containing peptides. J Immunol Methods 2005, 304 (1-2), 100–6.

