## Supplementary Information for "Exo-enzymatic addition of diazirine-modified sialic acid to cell surfaces enables photocrosslinking of glycoproteins"

Contents


| | |
|---|---|
| Page 2 | CMP-SiaDAz characterization |
| Page 3 | Figure S1. $^{1}$H NMR of CMP-SiaDAz |
| Page 4 | Figure S2. $^{13}$C NMR of CMP-SiaDAz |
| Page 5 | Figure S3. $^{31}$P NMR of CMP-SiaDAz |
| Page 6 | Figure S4. HPLC analysis of CMP-SiaDAz synthesis and purification |
| Page 7 | Figure S5. Treatment with *A. ureafaciens* neuraminidase removes α2-3Neu5Ac from Jurkat cell surfaces. |

**CMP-SiaDAz characterization**

$^{1}H$ NMR (600 MHz, $D_2O$) δ 7.93 (d, 1H, *J* = 7.6 Hz, H-6 of cytidine), 6.09 (d, 1H, J = 7.6 Hz, H-5 of cytidine), 5.96 (d, 1H , *J* = 4.5 Hz, H-1 of ribose), 4.33 – 4.30 (m, 2H, H-2 of ribose,H-3 of ribose), 4.28 – 4.26 (m, 3H, H-5a, H-5b,H-4 of ribose), 4.22 – 4.19 (m, 3H), 3.99 – 3.95 (m, 2H), 3.90 – 3.82 (m, 3H), 3.73 (ddd, *J* = 9.3, 6.7, 2.7 Hz, 1H), 3.59 – 3.53 (m, 2H), 2.46 (dd, *J* = 13.3, 4.7 Hz, 1H), 1.79 (dd, *J* = 12.9, 11.4 Hz, 2H), 1.70 (td, 1H), 1.00 (s, 3H). $^{13}C$ NMR (151 MHz, $D_2O$) δ 176.74, 175.84, 166.11, 157.74, 141.53, 96.34, 89.00, 82.85, 82.79, 74.18, 71.72, 70.33, 70.13, 68.56, 67.16, 63.30, 52.15, 39.39, 30.27, 29.53, 23.18, 18.51. $^{31}P$ NMR (243 MHz, $D_2O$) δ -4.70. HRMS: $C_{23}H_{34}N_6O_{16}P$(M-1), calcd 681.17744, found 681.17714.

**HPLC**

HPLC analysis was carried out using a SeQuant® ZIC®-HILIC (5µm, 200Å, 150 x10 mm) column. UV absorbance was monitored at 260 nm and the system was controlled via Chromeleon 7 chromatography software. Buffer A (acetonitrile), Buffer B (25 mM ammonium acetate) were used in gradient conditions for separation of compounds. Gradient condition was: 15 % buffer B for 3 min, 15–35 % buffer B over 30 min, 35–15 % buffer B over 2 min, 15 % buffer B for 10 min, followed by an equilibration phase of 90% buffer B for 3 min. Flow-rate was maintained at 3 mL/min for all separations.

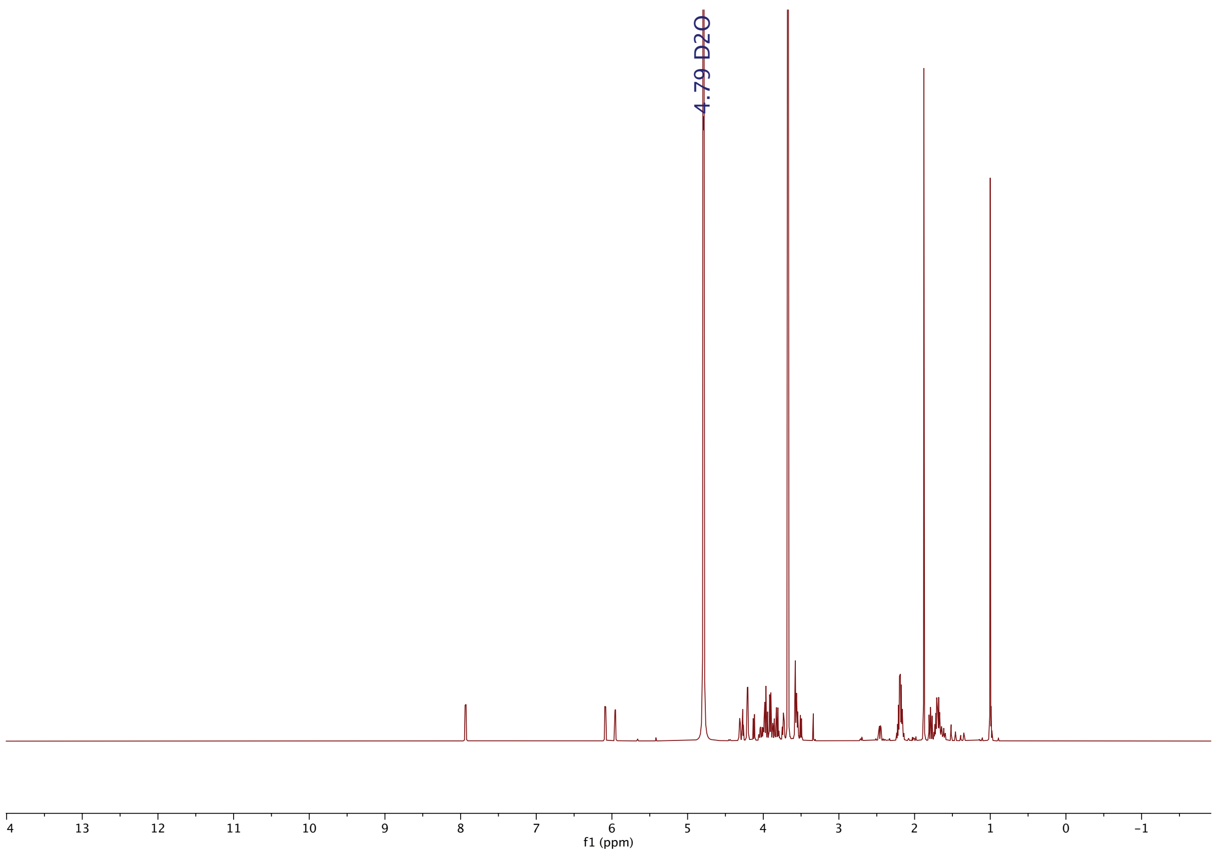


**Figure S1. $^{1}$H NMR of CMP-SiaDAz.**

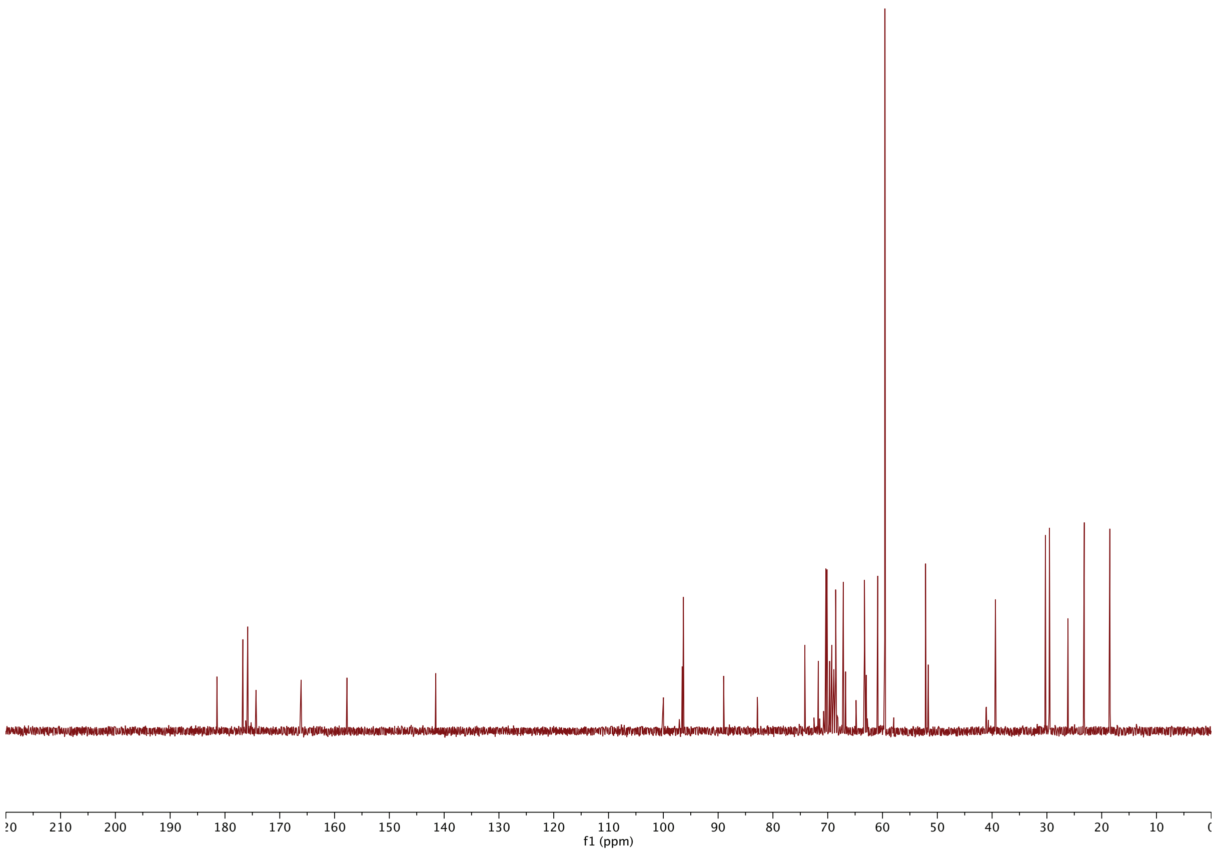


**Figure S2.** $^{13}$**C NMR of CMP-SiaDAz.**

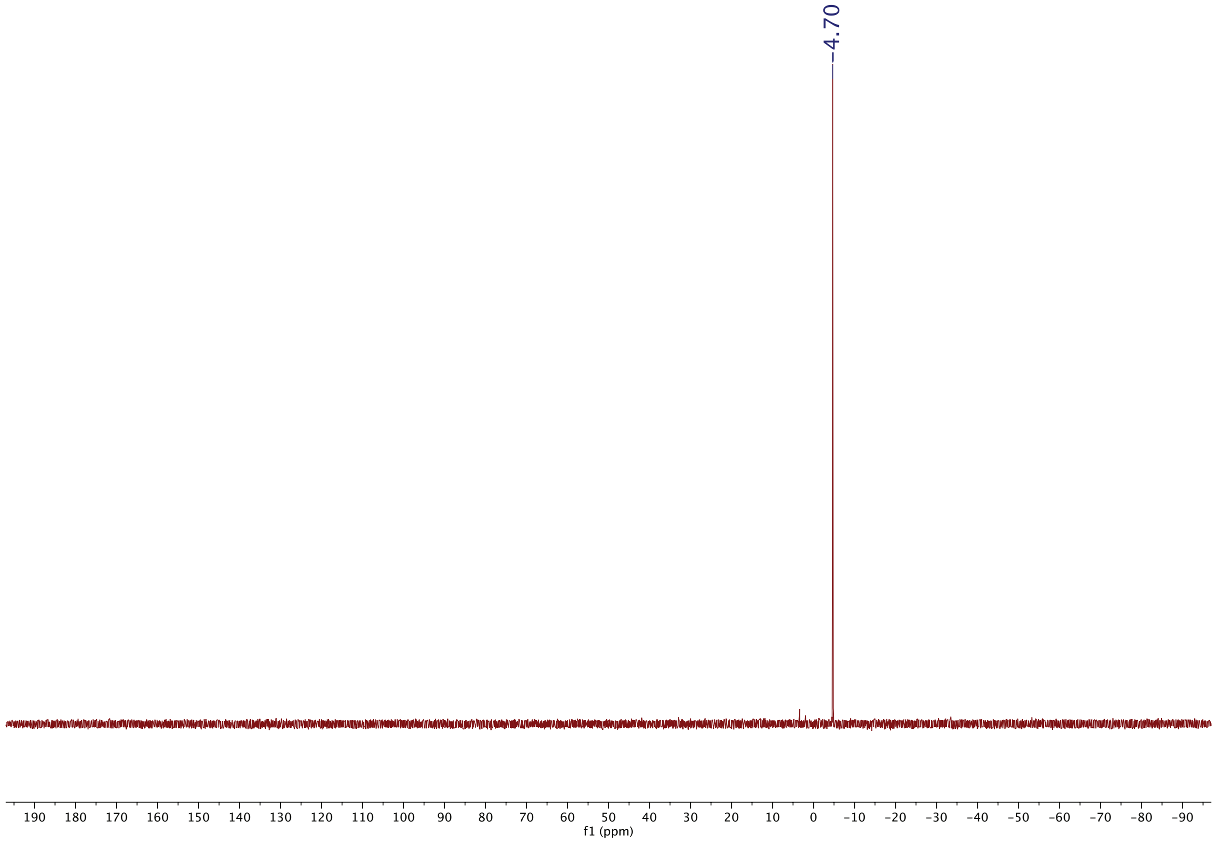


**Figure S3. $^{31}$P NMR of CMP-SiaDAz.**

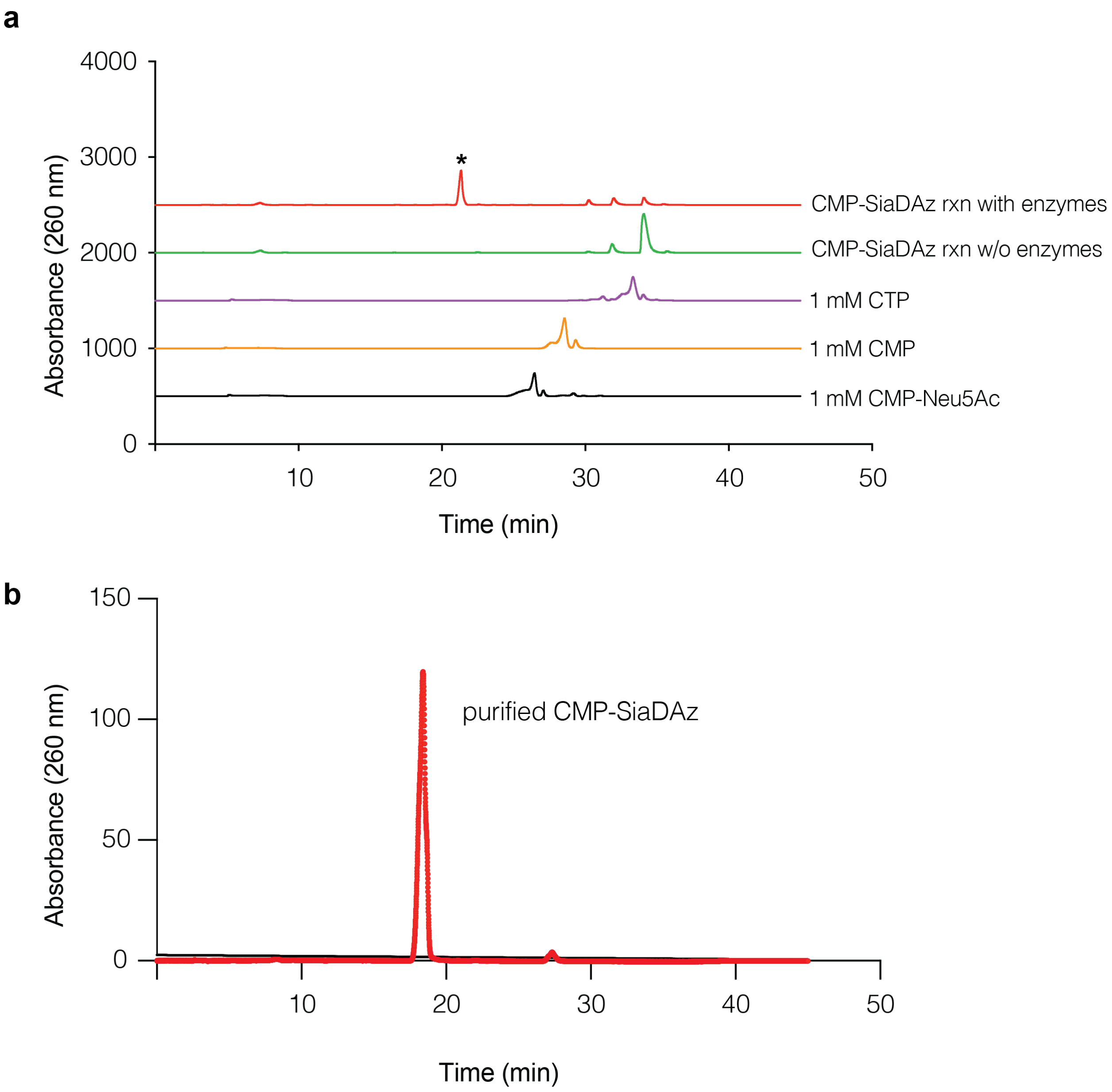


**Figure S4. HPLC analysis of CMP-SiaDAz synthesis and purification.** (**a**) HPLC chromatogram of crude CMP-SiaDAz reaction mixture, reactant mixture without aldolase or CMP-sialic acid synthase, and standards (CMP, CTP, and CMP-Neu5Ac). (**b**) HPLC chromatogram of CMP-SiaDAz purified using Biogel-P2 column.

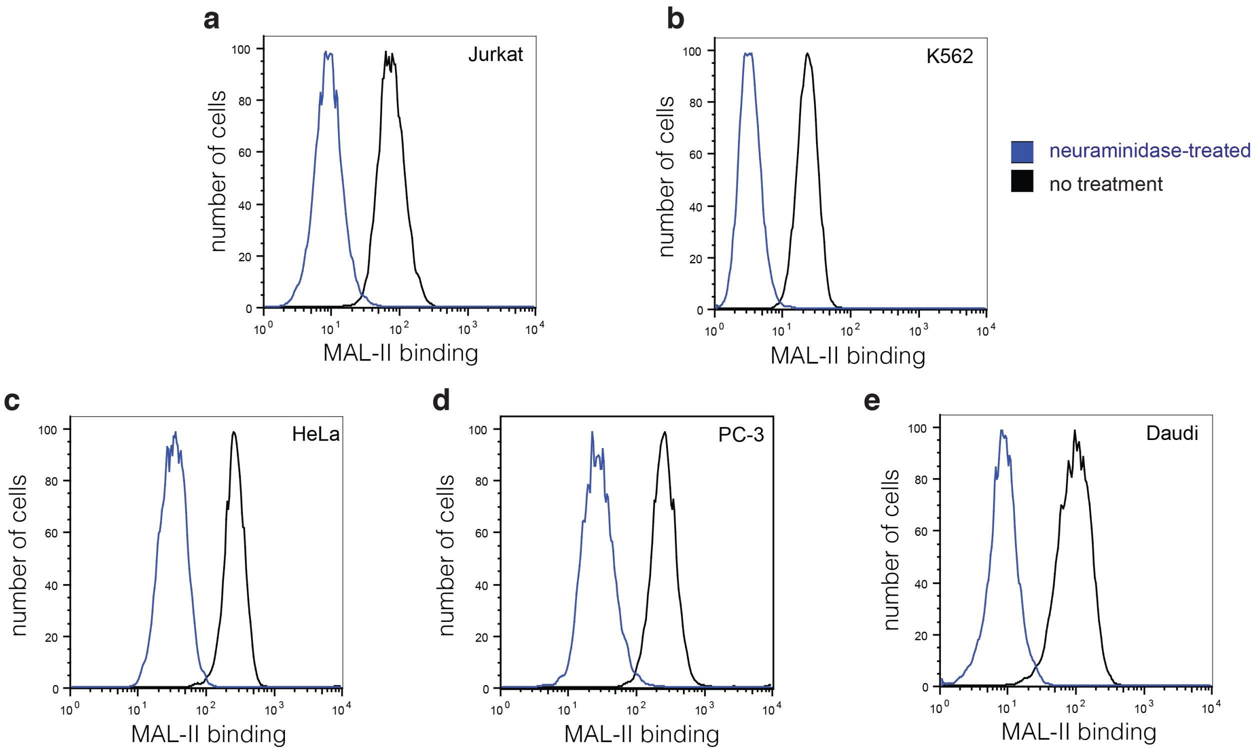


**Figure S5. Treatment with *A. ureafaciens* neuraminidase removes α2-3Neu5Ac from Jurkat cell surfaces.** Jurkat (**a**), K562 (**b**), HeLa (**c**), PC-3 (**d**), and Daudi (**e**) cells were treated with *A. ureafaciens* neuraminidase or not. Binding of MAL II-DTAF was measured by flow cytometry. Data presented are representative of three biological replicates.
